# Cyclin G2 inhibits oral squamous cell carcinoma growth and metastasis by binding to insulin-like growth factor binding protein 3 and regulating the FAK-SRC-STAT signaling pathway

**DOI:** 10.1101/2020.07.21.214361

**Authors:** Danning Wang, Jinlan Gao, Chenyang Zhao, Sen Li, Di Zhang, Xiaoyu Hou, Xinbin Zhuang, Qi Liu, Yang Luo

**Affiliations:** The Research Center for Medical Genomics, Key Laboratory of Cell Biology, Key Laboratory of Medical Cell Biology, Ministry of Education, School of Life Sciences, China Medical University, Shenyang, China

**Keywords:** cell cycle protein cyclin G2, OSCC, tumour suppressor, insulin-like growth factor binding protein 3, integrin, focal adhesion kinase, c-Src

## Abstract

The cell cycle protein cyclin G2 is considered a tumor suppressor. However, its regulatory effects and potential mechanisms in oral cancers are not well understood. This study aimed to investigate the effect of cyclin G2 on oral squamous cell carcinoma (OSCC). The data from 80 patients with OSCC were utilized to predict the abnormal expression of cyclin G2. The proliferation and metastasis were determined by a cell counting Kit-8 assay, flow cytometry, a wound-healing assay and a cell invasion assay. The expression of key proteins and genes associated with the cyclin G2 signaling pathways was determined by western blotting and real-time PCR, respectively. The orthotopic nude mice model was established by a mouth injection of SCC9 cells overexpressing cyclin G2. We showed that the low level of cyclin G2 in OSCC, which is negatively correlated with clinical staging, was a negative prognostic factor for the disease. We also found that cyclin G2 inhibited the proliferation, metastasis and blocked the cell cycle at G1/S of OSCC cells, suggesting that cyclin G2 has an inhibitory effect in OSCC. Mechanistically, cyclin G2 inhibited the growth and metastasis of OSCC by binding to insulin-like growth factor binding protein 3 (IGFBP3) and regulating the focal adhesion kinase (FAK) -SRC-STAT signal transduction pathway. Cyclin G2 competed with integrin to bind to IGFBP3; the binding between integrin and IGFBP3 was reduced after cyclin G2 overexpression, thereby inhibiting the phosphorylation of FAK and SRC. These results showed that cyclin G2 inhibited the progression of OSCC by interacting with IGFBP3 and that it may be a new target for OSCC treatment.

## Introduction

Oral squamous cell carcinoma (OSCC) is the sixth most common malignant tumour in the world and the most common malignant oral cancer tumor, accounting for more than 90% of oral cancer cases. More than 300,000 new cases are diagnosed each year, and the morbidity and mortality among young people have continually risen in recent years (1, 2). Despite recent advances in radiotherapy, chemotherapy and traditional surgery, the 5-year survival rate for OSCC is still only about 50% (3). Although the cure rate of early OSCC exceeds 80%, more than 70% of patients with advanced OSCC cannot be cured (4, 5). It is generally believed that OSCC has a high potential for local invasion and lymph node metastasis, and clinical staging plays an important role in the survival prediction of OSCC patients (6). Nonetheless, the precise mechanisms leading to OSCC are not fully understood.

Cyclin G2 is an unconventional cell cycle protein encoded by the *CCNG2* gene that plays a negative role in the cell cycle process (7). Thus, it also plays an important role in the biological functions of cell growth inhibition and tumor suppression (8). Cyclin G2 is a potential cancer suppressor gene whose expression may be associated with the pathological process underlying tumor development (9). The expression levels of cyclin G2 are reduced in thyroid, breast, kidney, stomach, esophageal and other tumors according to early reports (10–14). However, there are few reports on the identity of pathways and the precise mechanisms that mediate the roles of cyclin G2 in OSCC and other cancers.

Thus, the goal of our research was to explore the relationship between cyclin G2 and the growth and metastasis in OSCC and the underlying molecular mechanisms. Through a mass spectrometric analysis, we identified that IGFBP3 was a possible cyclin G2 interacting protein. IGFBP3 promotes OSCC cell migration and lymph node metastasis through integrin β1 in an insulin-independent manner (15). Inside-out signaling through the integrin-FAK axis may regulate cancer cell adhesion, proliferation and metastasis (16). Furthermore, we found that the combination between integrin and IGFBP3, the phosphorylation of FAK and SRC and the FAK-SRC-STAT axis were negatively regulated after cyclin G2 overexpression. Taken together, these results indicated that cyclin G2 functioned as a tumor suppressor in OSCC through its interaction with IGFBP3 and the subsequent blockage of the connection between integrin and IGFBP3, the phosphorylation of FAK and the FAK-SRC-STAT signaling pathways. When the phosphorylation of STAT3 was reduced, its translocation to the nucleus was reduced, which decreased the transcriptional regulation of *Bcl-2, c-Myc,* and *MMP9,* thereby regulating cell growth and metastasis. STAT3 is constitutively active in a variety of cancers. This activation deregulates the signaling that controls cell proliferation, including signals involving Bcl-2 and c-Myc, and the epithelial to mesenchymal transition, which involves MMP-9. This deregulation promotes tumor progression (17). This study demonstrated the inhibitory function of cyclin G2 in OSCC growth and metastasis and explored the underlying mechanisms. Thus, Cyclin G2 may be a key treatment target of OSCC.

## Methods

### Patients and tissue

The tissue samples were collected from 80 patients diagnosed with OSCC who received treatment at the Oral Hospital of China Medical University between January 2018 and March 2019. The patients underwent extensive OSCC resection during the surgery, and OSCC tissue and paired normal tissue were obtained for postoperative analysis. In addition, clinical data were collected, including the patients’ age and sex, tumor size, metastasis status and clinical stage.

### Hematoxylin and eosin staining and immunohistochemistry

The OSCC tissues from each group were paraffin embedded and were cut on a microtome into 4 μm sections. The tissues were stained using a hematoxylin-eosin staining kit (Wanlei, Shenyang, China) and were observed under a microscope (Olympus, Tokyo, Japan). In addition, some of the tissue sections were dewaxed in xylene, rehydrated with alcohol, treated for antigen-retrieval, incubated in hydrogen peroxide, blocked with goat serum and then were incubated with one of the following primary antibodies: anti-cyclin G2 antibody (HPA034684, Sigma-Aldrich, Santa Clara, CA, USA); phospho-FAK (ab81298, Tyr397, Abcam, Cambridge, MA, USA); phospho-SRC (AF3162, Tyr419, Affinity Biosciences, OH, USA) or phospho-STAT3 (AF3293, Tyr705, Affinity) overnight at 4°C. The membranes were incubated with an HRP-linked secondary antibody (Goat anti-Rabbit IgG (H+L), 31460, Thermo Fisher, Waltham, MA, USA), DAB stained, hematoxylin counter stained and were imaged under the microscope. A staining index (0–12) was determined by multiplying the score of the staining intensity by the score for the positive cells. The intensity was scored as follows: 0, negative; 1, weak; 2, moderate and 3, strong. The positive cell frequency was scored as follows: 0, less than 5%; 1, 5%–25%; 2, 26%–50%; 3, 51%–75% and 4, greater than 75%.

### Cell culture

The human OSCC cell line SCC-9 was purchased from Cobioer (Nanjing, China), and the cells were cultured in DMEM/F12 medium (including 10% fetal bovine serum and 0.4 μg/ml hydrocortisone). The cells were maintained in an incubator at 37°C and with 5% CO_2_.

### Cell infection

To generate cell lines that stably overexpressed cyclin G2, FLAG-tagged cyclin G2 (FLAG-*CCNG2*) was cloned into a lentivirus-GFP lentiviral vector and was packaged by GeneChem Co., Ltd (Shanghai, China) used as previously described (18). A GFP-lentiviral vector was used as a negative control. The SCC-9 cells were plated at a density of 1 × 10^5^ cells per well in 24-well plates. When the cell confluence reached 70%, the cells were infected with the cyclin G2 overexpressing and negative control lentiviral vectors. The transduced cells were selected with puromycin.

### Cell proliferation assay

The cells were plated in 96-well plates at a density of 5 × 10^3^ cells per well for 0 h, 24 h, 48 h and 72 h. CCK8 (Beyotime Biotechnology, Shanghai, China) was added to each well, and the plates were incubated for 2 hours at 37°C. The absorbance was measured at 490 nm using an ultraviolet spectrophotometer (Thermo Fisher Scientific, Waltham, MA, USA).

### Cell cycle detection

A total of 5 × 10^5^ cells was collected and fixed in 70% ethanol overnight and were then incubated with a propidium iodide stain (BD Biosciences, Franklin Lakes, NJ, USA) at 4°C. The percentage of cells in the different cell cycles was then determined.

### Wound-healing assay

The cells were plated at a density of 5 × 10^5^ per well in a 6-well plate. Then, a 2-mm wide plastic pipette tip was used to scrap the cells to perform wound healing assays. The cells were observed and imaged under a microscope at 0 h and 24 h. The wound-healing rate was calculated as (the 0 h cell wound width – the experimental point cell wound width)/0 h cell wound width ×100%.

### Cell invasion assay

The chambers (8μm pore size, corning, NY, USA) were pre-coated with Matrigel (BD Biosciences, NJ, USA). The cell concentration was adjusted to 5 × 10^4^/ml in serum-free medium, and 200 μl of cell suspension was seeded to the upper chamber; complete culture medium was added to the lower layer. After 24 h in culture, the upper matrix material was removed using a cotton swab, and the cells were fixed and stained with crystal violet. Then, 5 randomly chosen fields were counted to determine the cell migration indices, and these were imaged under a microscope.

### Production of MMP9

The cells were plated at a density of 5 × 10^4^ per well in 24-well plates. The supernatant was collected after the 24-h and 48-h culture, and MMP9 levels were determined using an MMP9 ELISA kit (Boster Biosciences, Wuhan, China). The methods were as described in the manufacturer’s protocol. The supernatants were collected, and the MMP9 concentrations were measured at 450 nm. The values were calculated using a standard curve.

### Mass spectrometry

The mass spectrometry was performed according to the methods reported in our previous study (19). The cells expressed FLAG or FLAG-tagged cyclin G2 was lysed using RIPA lysis buffer. The immunoprecipitation assay was performed to separate the FLAG-tagged protein from the lysate using the anti-FLAG affinity gel. After the anti-FLAG affinity gel was washed in PBS and resuspended in 2×loading buffer, the proteins were subjected to polyacrylamide gel and stained with Coomassie Brilliant Blue (R250). The aim bands were harvested and digested to conduct mass spectrometry.

### Co-immunoprecipitation and Western blot

The cell lysates were extracted with RIPA lysis buffer that contained a protease inhibitor cocktail and phosphatase inhibitor cocktail (Roche, Basel, Switzerland). Equivalent concentrations of protein were separated using gel electrophoresis and were then transferred to a PVDF membrane. The membranes were incubated with one of the following primary antibodies: anti-cyclin G2 (1:500, HPA034684, Sigma); Flag (1:500, ab49763, Abcam); Integrin β1 (1:1000, ab52971, Abcam); FAK (1:1000, bs-1340R, Bioss, Beijing, China); phospho-FAK (1:1000, ab81298, Tyr397, Abcam); SRC (1:500, WL01570, Wanlei); phospho-SRC (1:1000, AF3162, Tyr419, Affinity); STAT3 (1:300, WL01836, Wanlei); phospho-STAT3 (1:1000, AF3294, Affinity); Bcl-2 (1:1000, ab32124, Abcam); c-Myc (1:500, 9E10, Santa Cruz Biotechnology, CA, USA); MMP9 (1:500, 10375-2-AP, Proteintech, ID, USA); β-actin (1:1000, ab6276, Abcam) and GAPDH (1:1000, 5174, Cell Signaling Technology, Danvers, MA, USA). The protein levels were determined using an ECL Plus kit.

### RT-qPCR

After cell lysis, TRIzol (Invitrogen) was added to extract the RNA, and high-fidelity enzymes were used to convert it into cDNA. The following primers were designed: CCNG2 F: 5‘ CGGAGAATGATAACACTTTG, *CCNG2* R: 5‘ GTTTCACCTTCATAAGAGCC; *IGFBP3* F: 5‘ AAATGCTAGTGAGTCGGAGGAA, *IGFBP3* R: 5‘ GATGATTATCTTTGAATGGAGGG; *c-Myc* F: 5‘ CACCCTTCTCCCTTCGG, *c-Myc* R: 5‘ CAGTCCTGGATGATGATGTTT; *Bcl2* F: 5‘ TGTGGCCTTCTTTGAGTTCG, *Bcl-2* R: 5‘ CATCCCAGCCTCCGTTATCC; *MMP9* F: 5‘ TTTGACAGCGACAAGAAGTG, *MMP9* R: 5‘ CAGGGCCGAGGACCATAGAGG and *GAPDH* F: 5‘ TGTTGCCATCAATGACCCCTT, *GAPDH* R: 5‘ TCCACGACGTACTCAGCG. The primers were synthesized by Sangon Biotech (Shanghai, China).

### Animal experiments

BALB/C nude mice (5- to 6-week-old females, 18–20 g) were purchased from Beijing Vital River Laboratory Animal Technology Co., Ltd. The mice were randomly divided into two groups (n = 6/ group). After abdominal cavity anesthesia with 1% sodium pentobarbital at a concentration of 500 mg/kg, a total of 1 × 10^7^ SCC9 cells overexpressing cyclin G2 or control GFP (NC) cells in 50μl of PBS was injected into the tongue of the nude mice (20). Mice were housed at room with 40-60% humidity, and with a light cycle of 12 hour light and12 hour dark under pathogen-free conditions. Animal health and behaviour were monitored every day. All of the mice were maintained using a homemade semi-flow feeding system, which refers to the recipe of rodent diet (21) and contained 35% ewe’s milk, 35% corn starch, 10% gelatin, 10% eggs, 5% corn oil and a 5% sugar and vitamin mix. One mouse lost weight and starved to death on the 20th day, so 12 mice (6 pairs) were included in the end. All of the twelve mice were euthanized via anesthesia overdose in 42 days after injection (22), and the tumors were isolated. We then measured the weight and volume using the formula V=length*width*height and also extracted the swollen lymph nodes, followed by fixing and embedding.

### Statistical analysis

All the experiments were repeated at least three times, and the results are expressed as the mean ± standard deviation. SPSS 19.0 software was used for the t-test, chi-square test and ANOVA. P < 0.05 was considered to indicate statistical significance.

## Results

### Cyclin G2 expression is decreased in human OSCC tissue

Immunohistochemistry was performed on 80 OSCC tissue samples and 10 adjacent normal tissue samples to assess cyclin G2 expression, and the results were analyzed in combination with the clinical data. Age, sex and tumor size were not correlated with the expression of cyclin G2, whereas the clinical stage was associated with cyclin G2 expression (P<0.05). The expression of cyclin G2 was significantly reduced in the stage III and IV samples and was negatively correlated with the clinical stage (**Table I**, **Figs. 1A and B**).

**Figure 1.**
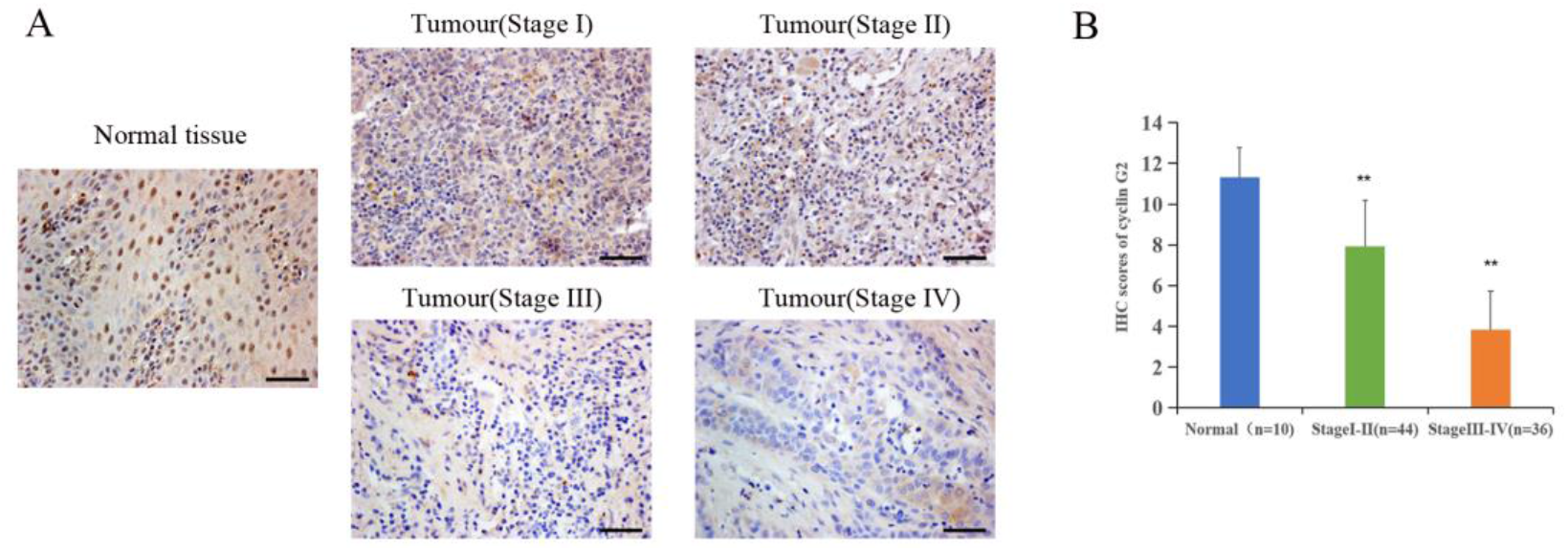
Cyclin G2 expression negatively correlates with clinical stage in OSCC. A: The expression of cyclin G2 in OSCC at the different clinical stages by an immunohistochemical analysis. Scale bar = 100 μm. B Immunohistochemical scores of cyclin G2 at the different clinical stages. \**p*<0.05 and *\*\*p* <0.01 vs. Normal group.

**Table1.**
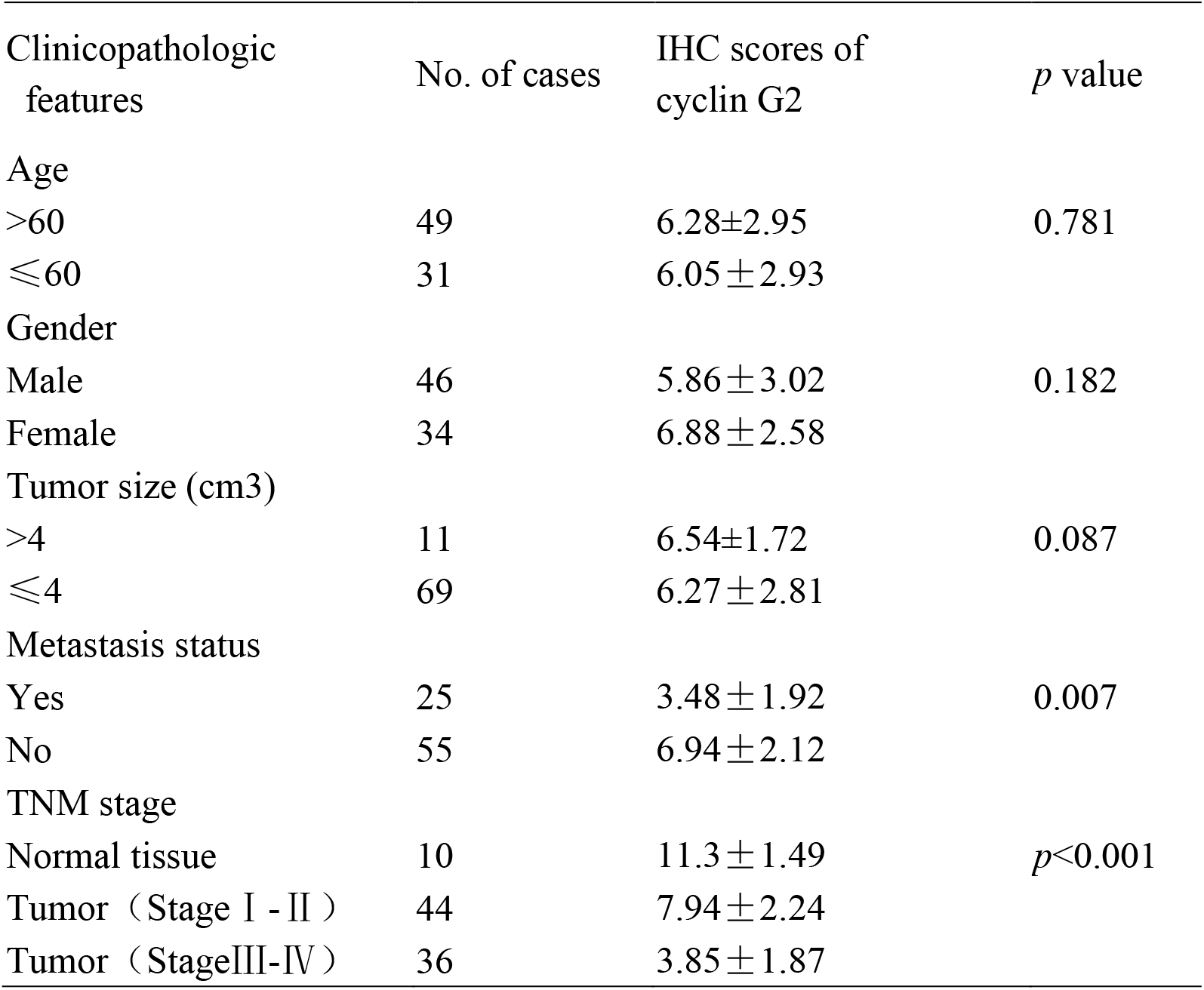
Relationship between positive immunohistochemical expression of cyclin G2 and the clinical characteristics of oral squamous cancer

| Clinicopathologic features | No. of cases | IHC scores of cyclin G2 | *p* value |
| --- | --- | --- | --- |
| Age |  |  |  |
| >60 | 49 | 6.28±2.95 | 0.781 |
| ≤60 | 31 | 6.05±2.93 |  |
| Gender |  |  |  |
| Male | 46 | 5.86±3.02 | 0.182 |
| Female | 34 | 6.88±2.58 |  |
| Tumor size (cm3) |  |  |  |
| >4 | 11 | 6.54±1.72 | 0.087 |
| ≤4 | 69 | 6.27±2.81 |  |
| Metastasis status |  |  |  |
| Yes | 25 | 3.48±1.92 | 0.007 |
| No | 55 | 6.94±2.12 |  |
| TNM stage |  |  |  |
| Normal tissue | 10 | 11.3±1.49 | *p*<0.001 |
| Tumor (Stage I - II ) | 44 | 7.94±2.24 |  |
| Tumor (StageIII-IV ) | 36 | 3.85±1.87 |  |

### Cyclin G2 inhibits the proliferation and metastasis of OSCC cells and blocks the cell cycle at G1/S

To study the effect of cyclin G2 on the proliferation of OSCC, we first constructed stably transfected SCC-9 cell lines that overexpressed cyclin G2 and control cells that had GFP overexpression (negative control, NC). The CCK8 assay showed that cell proliferation was significantly inhibited in the SCC-9 cells after cyclin G2 overexpression (**Fig. 2A**). Then, the effect of the overexpression of cyclin G2 on the cell cycle was tested in the SCC-9 cells using flow cytometry. The results showed that the number of G1 cells increased and the number of S cells decreased with cyclin G2 overexpression (**Figs. 2B and C**). Thus, cyclin G2 blocks the cell cycle in OSCC cells at the G1/S phase.

**Figure 2.**
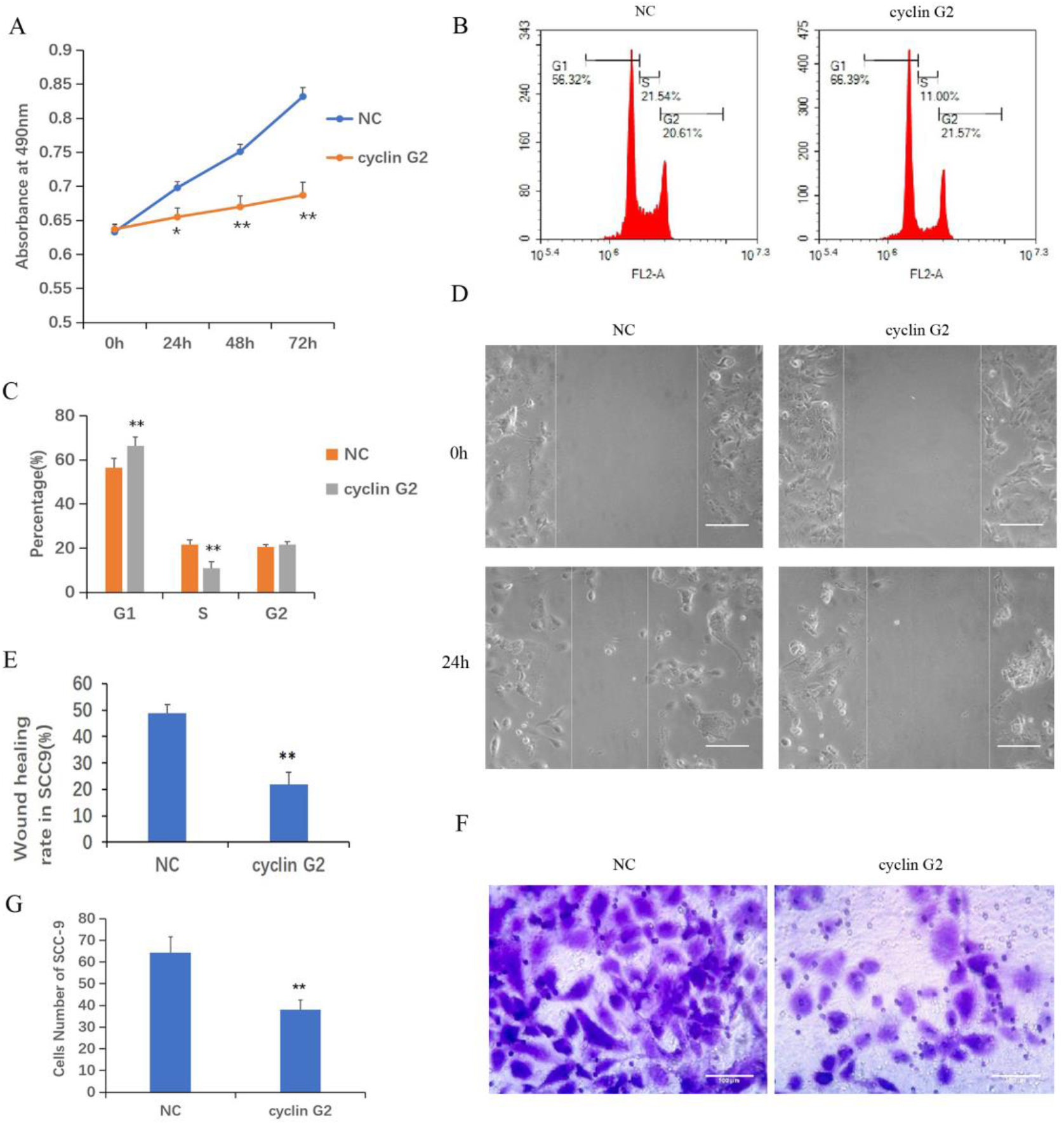
Cyclin G2 inhibits the proliferation and metastasis of OSCC cells and blocks the cell cycle at G1/S. A: Absorbance of the SCC-9 cells in the CCK8 cell proliferation experiment. B and C: Flow cytometry detection of the cell cycle in the SCC-9 cells. D and E: Representative photos and quantification of the wounded areas after wounding the SCC-9 cells. Scale bar =100 μm. F and G: Analysis of the invasion and quantification of the SCC-9 cells. Scale bar = 100 μm. *p < 0.05, **p < 0.01 vs. vector.

To clarify the effects of cyclin G2 on OSCC migration and invasion, we tested the effect of cyclin G2 overexpression in the SCC-9 cells using the wound healing assay and Matrigel transwell assay. The wound-healing rates of the SCC-9 cells in the NC and cyclin G2 groups were 48.67%±3.45% and 21.89%±4.57%, respectively. Thus, the wound healing capacity of the SCC-9 cells was significantly inhibited in the cyclin G2 overexpression group. In addition, cell migration was significantly reduced after cyclin G2 overexpression (**Fig.2D and E**). In the Matrigel transwell experiment, the number of cells passing through the Matrigel and substrate membrane was counted, and the SCC-9 cell numbers for the NC and cyclin G2 groups were 64.2±7.3 and 38.0±4.5, respectively. In summary, cyclin G2 overexpression inhibits the migration and invasion abilities of the SCC-9 cells (**Figs. 2F and G**).

### Cyclin G2 inhibits the growth and metastasis of OSCC *in vivo*

The nude mice were maintained for 42 days after the injection of the SCC9 cells. During that time, tumors formed in the rear of the tongue and the lymph nodes became swollen (**Fig. 3A**). The results showed that the volume and weight of the tumors were significantly reduced by the overexpression of cyclin G2 (**Figs. 3B-D**). The tumors in the mouth and the swollen lymph nodes were stained with hematoxylin and eosin. OSCC formed in both the cyclin G2 overexpression group and the negative control group. However, the pathological differentiation degree of the tumors was higher in the cyclin G2 overexpression group. The swollen lymph nodes in the cyclin G2 group showed just inflammation, whereas the lymph nodes in the control group showed a significant level of squamous cancer metastasis. We also assessed Ki67 in the two groups of tumors and lymph nodes, and the Ki67 expression was significantly lower in the cyclin G2 group than in the control group (**Figs. 3E and F**).

**Figure 3.**
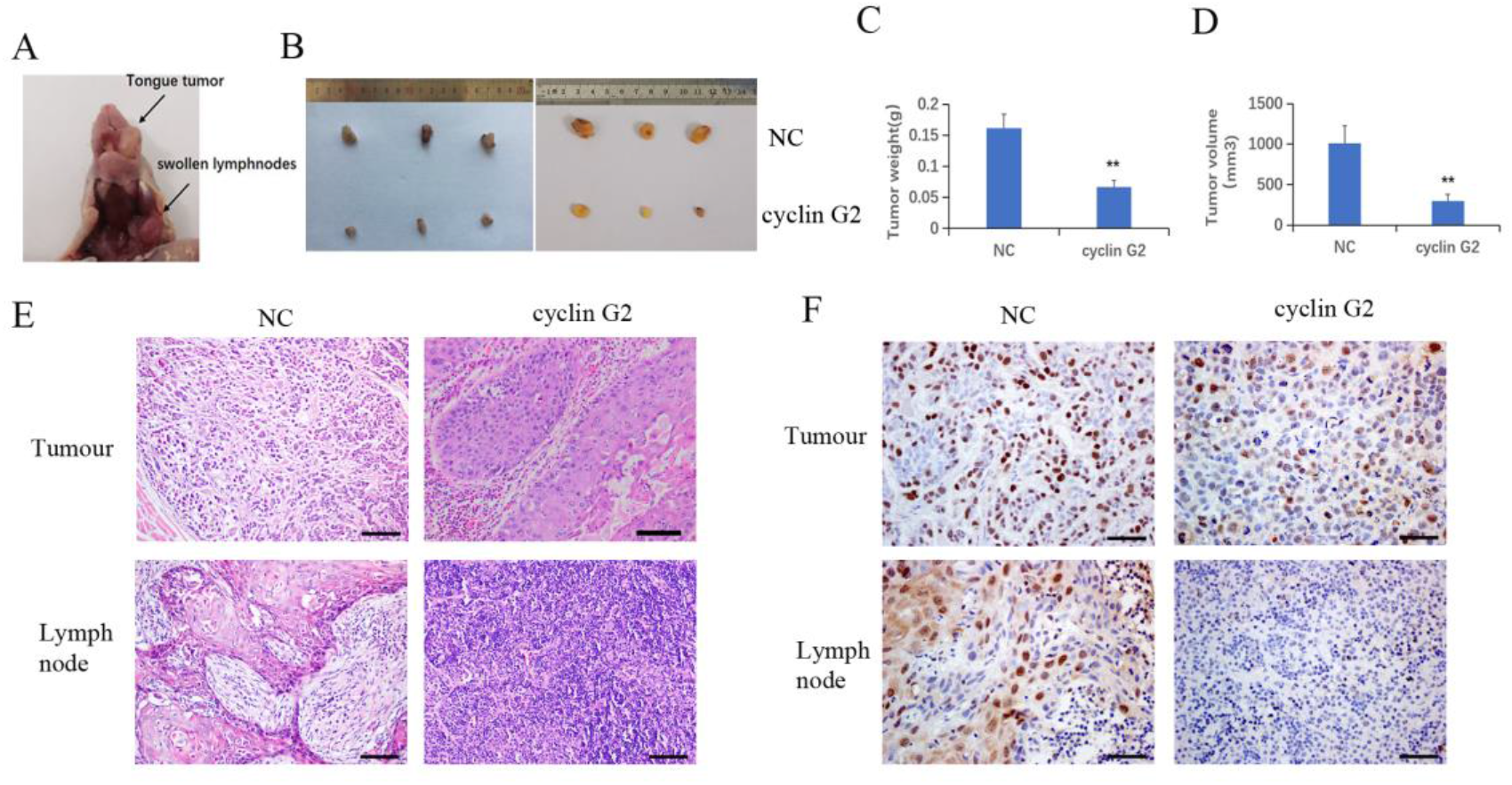
Cyclin G2 inhibits the growth and metastasis of OSCC *in vivo.* A: The tumor in the rear of the tongue and the swollen lymph nodes after 42 days. B: Overall images of the tumors and lymph nodes. C and D: Tumor volumes and weights in the cyclin G2-overexpressing and negative control groups. E: Hematoxylin and eosin staining of the tumor in the rear of the tongue and the swollen lymph nodes. F: Immunohistochemical staining of Ki67 in the oral tumors and lymph nodes. Scale bar = 100 μm. *p < 0.05, **p < 0.01 vs. vector.

### Cyclin G2 inhibits the binding between IGFBP3 and integrin by interacting with IGFBP3

To study the molecular mechanism underlying the ability of cyclin G2 to inhibit growth and metastasis in OSCC, we looked for potential protein interactions with cyclin G2 using mass spectroscopy experiments. After screening, we identified IGFBP3 as a possible protein interacting with cyclin G2. We previously published that cyclin G2 also interacted with Dapper 1 and lactate dehydrogenase A, which both further supported the role of cyclin G2 as tumor suppression (18, 19). To validate the findings here, we first extracted proteins from cyclin G2 overexpressing SCC-9 cells, and through the Co-IP experiment, we found that cyclin G2 did indeed interact with IGFBP3 (**Fig. 4A**). Next, we performed an immunobifluorescence analysis of cyclin G2 and IGFBP3 in the SCC-9 cells, and cyclin G2 and IGFBP3 were co-located in the cytoplasm (**Fig. 4B**). IGFBP3 promotes cell migration and lymph node metastasis in OSCC cells by the requirement of integrin β1 (15). IGFBP-3 binds to the β1-integrin subunits and manipulates the activity of integrin and the integrin receptor complexes and the downstream intracellular signaling pathways (23, 24). However, we found that these combination effects were weakened between IGFBP3 and integrin after cyclin G2 overexpression in the SCC9 cells (**Fig. 4C**). This demonstrated that the activation effect of IGFBP3 in integrin was inhibited after cyclin G2 overexpression. The integrin receptor complex was inhibited because of the reduced activity of integrin. Additionally, the formation of integrin receptors complexes was recognized to be the critical determinant of signaling from integrin receptors, and thus, the downstream intracellular signaling pathways maybe affected.

**Figure 4.**
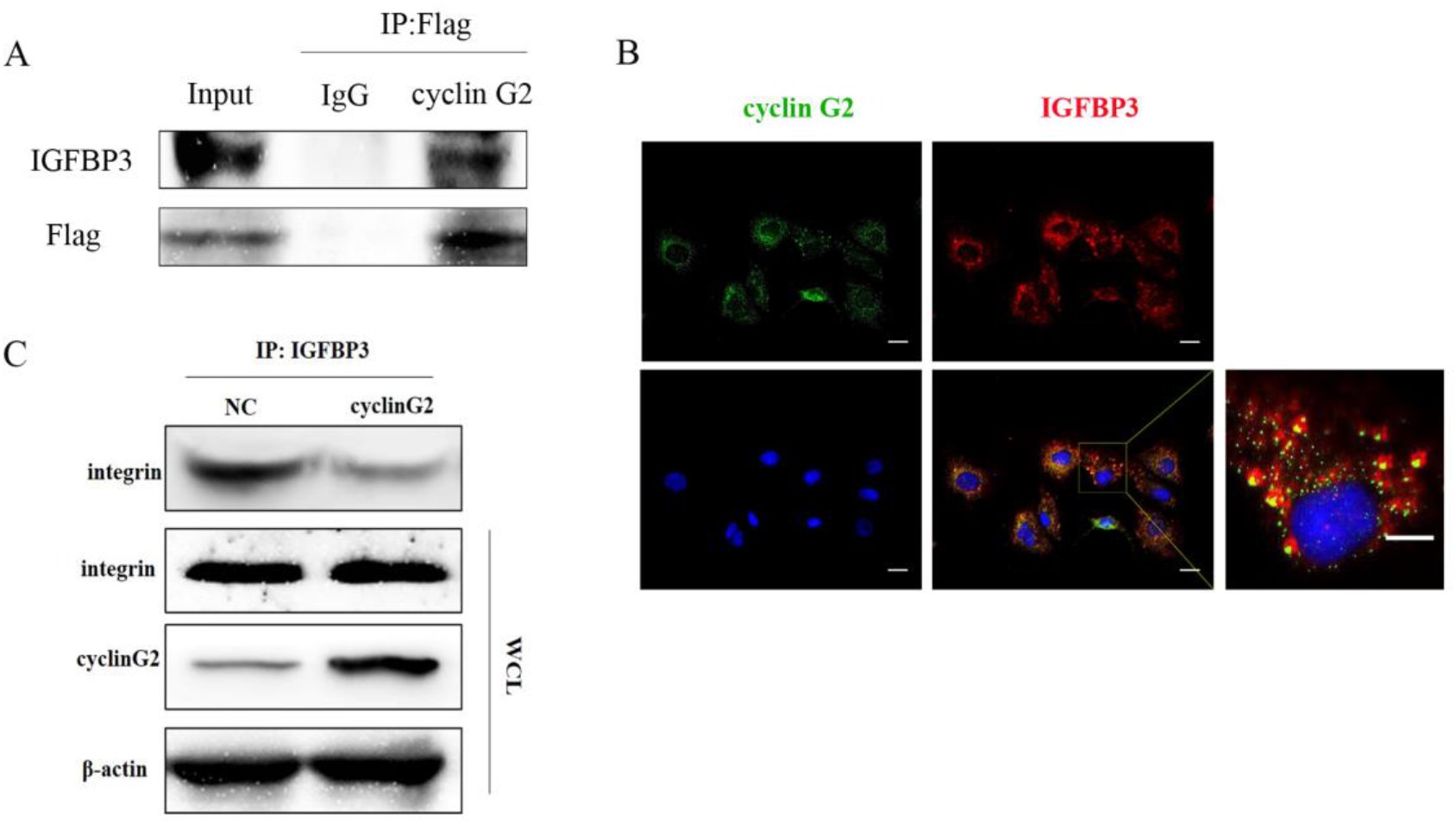
Cyclin G2 inhibits the binding between IGFBP3 and integrin by interacting with IGFBP3. A: Co-IP analysis of the interaction between cyclin G2 and IGFBP3 in SCC-9 cells. B: Cyclin G2 and IGFBP3 co-localized in the cytoplasm of the SCC-9 cells. C: The combination effects of IGFBP3 and integrin were reduced after cyclin G2 overexpression.

### Cyclin G2 inhibits the FAK-SRC-STAT pathway *in vitro and in vivo*

IGFBP-3 modulates the classical integrin-mediated membrane recruitment of FAK and the subsequent activation of SRC (24). We found that the phosphorylation of FAK in the integrin pathway was inhibited after cyclin G2 overexpression in the SCC-9 cells, which inhibited the phosphorylation and activation of the downstream SRC-STAT pathway. STAT3 translocates to the nucleus after its phosphorylation, in order to regulate transcription. We detected the protein expression of the downstream genes *Bcl-2*, *c-Myc* and *MMP9*, and Bcl-2, c-Myc and MMP9 were decreased after cyclin G2 overexpression (**Fig. 5A**). In addition, RT-qPCR revealed that the mRNA levels of *Bcl-2*, *c-Myc* and *MMP9* decreased after cyclin G2 overexpression (**Figs. 5B-D**). Furthermore, we detected MMP9 secretion using an ELISA test. MMP9 secretion was reduced after cyclin G2 overexpression (**Fig. 5E**). Finally, the expression of cyclin G2, p-FAK, p-SRC and p-STAT3 *in vivo* were lower in the cyclin G2 overexpression group (**Figs. 5F and G**). Taken together, we concluded that cyclin G2 inhibited the growth and metastasis of OSCC via its interaction with IGFBP3.

**Figure 5.**
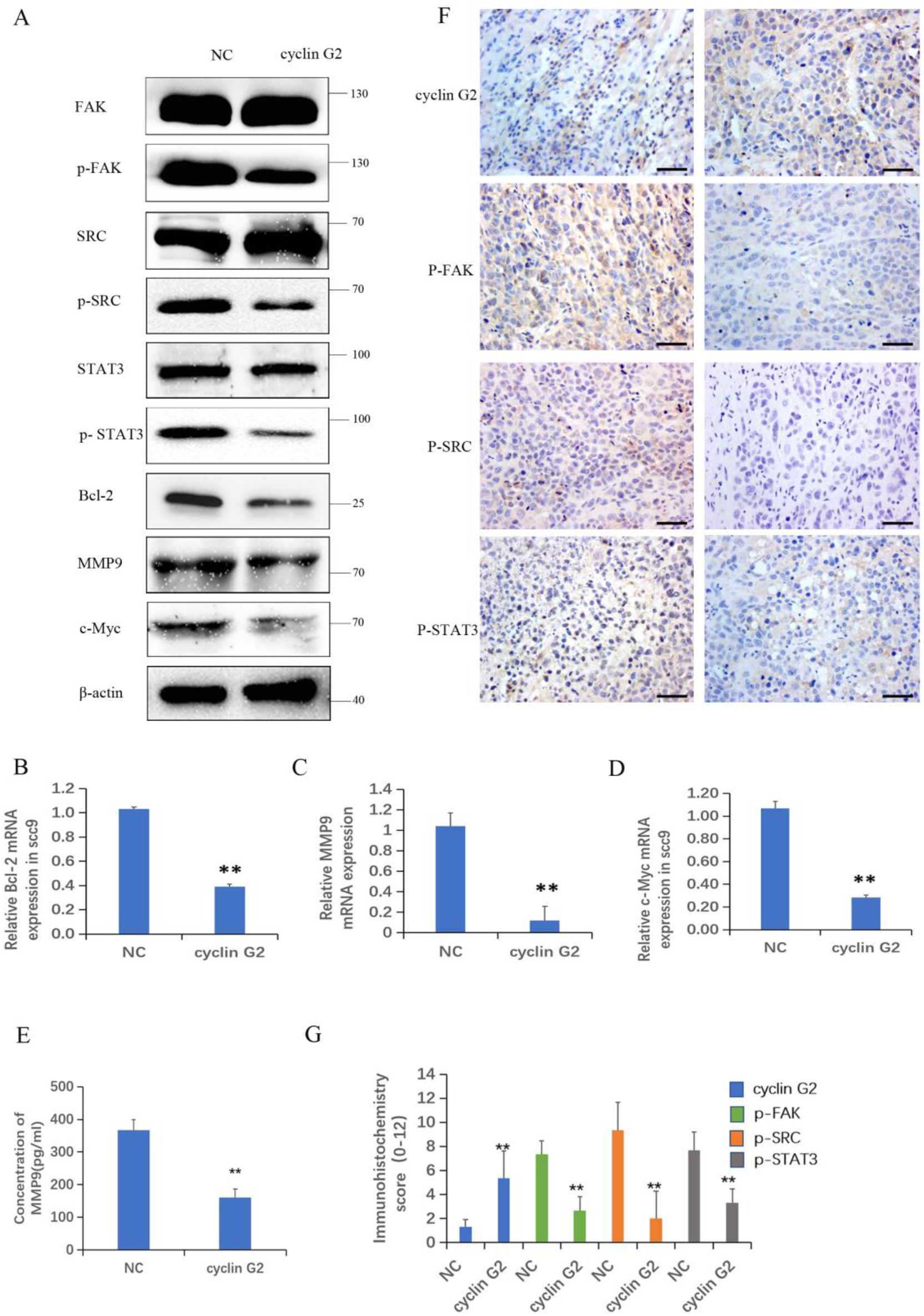
Cyclin G2 inhibits the FAK-SRC-STAT pathway *in vitro* and *in vivo.* A: The protein expression of FAK, P-FAK, SRC, P-SRC, STAT3, P-STAT3, Bcl-2, c-Myc and MMP9 in the FAK-SRC-STAT pathway. B-D: The relative mRNA expression of Bcl-2, c-Myc and MMP9. E: Analysis of MMP9 secretion and its quantification in SCC-9 cells. F and G: Immunohistochemical staining and analysis of the expression of cyclin G2, p-FAK, p-SRC and p-STAT3. Scale bar = 100 μm. *p < 0.05, **p < 0.01 vs. vector.

## Discussion

Cyclin G2 is a tumor suppressor and has a low expression in several cancers (25). In this study, we found that cyclin G2 levels were reduced in clinical OSCC tissue samples and that its expression was negatively correlated with the clinical stage of OSCC. The overexpression of cyclin G2 inhibited the proliferation and metastasis of OSCC cells both *in vitro* and *in vivo.* These findings suggest that cyclin G2 is an inhibitor of OSCC.

The current study demonstrated that the cyclin G2 interacts with IGFBP3. The protein is closely related to the development and metastasis of tumors. In addition, IGFBP3 not only enhances the growth of insulin-like growth factors (IGFS) (26), but also binds to the integrin β1 subunit in an insulin-independent manner. Also, it promotes the formation of a link between the integrin α and β subunits, thereby activating the integrin (27). The integrin receptor regulates the intracellular signaling pathways and the adhesion process of the cells and the extracellular matrix (28). Integrins connect to the actin cytoskeleton and transmit signals from the outside to the inside through focal adhesion multiprotein complexes. Stimulated integrins lead to the activation of the focal adhesion tyrosine kinases, FAK and SRC (29), which, in turn, promotes the activation of numerous downstream signaling pathways that play a pivotal role in cellular processes, such as adhesion, proliferation, migration and survival (30). Nuclear FAK is specifically linked to IGFBP3 through its interaction with Runx1, which leads to cell cycle progression and tumor growth regulation (31). IGFBP3 is overexpressed in a variety of tumors, but its biological significance remains unclear (32–35). The overexpression of IGFBP3 in OSCC promotes cell migration and lymph node metastasis in a manner that depends on integrin beta-1 (36).

Here, we revealed that cyclin G2 competed with integrin to bind to IGFBP3, and the binding between integrin and IGFBP3 was reduced after cyclin G2 overexpression, which impeded the formation of the integrin receptor complex, thereby inhibiting the phosphorylation of FAK and SRC. The Mbd domain of IGFBP3 (mature protein amino acid residues 215–241) interacts with integrin (26). A future study would be aimed to analyze whether cyclin G2 interacts with a part of the Mbd domain of IGFBP3 and further confirm the competitive role of cyclin G2 and integrin.

FAK is a non-receptor tyrosine kinase and is the central molecule in the transfer process of the integrin-mediated signaling effectuated via the Y397 phosphorylation of FAK (37). FAK increases the activity of SRC, which then catalyzes the tyrosine phosphorylation in FAK. Thus, the two proteins are mutually activated (38, 39). The FAK-SRC complex regulates cell growth, differentiation, metastasis and survival (40). The abnormally activated FAK and SRC proteins are associated with the progression of several types of tumors, including colon cancer, breast cancer, prostate cancer, lung cancer and oral cancer (41–45). Cyclin G2 inhibits the phosphorylation and activation of FAK and SRC, which affects the activation of downstream STAT3 and inhibits the signal transmission of the FAK-SRC-STAT axis. The regulation of the proliferation, invasion and metastasis of tumor cells allows for the interaction of phosphorylated FAK and SRC with the STAT3 signaling pathway. STAT3 is a transcription factor that is translocated to the nucleus via phosphorylation, thereby regulating an array of genes and the expression of proteins associated with growth and metastasis. When the phosphorylation of STAT3 is reduced, its translocation to the nucleus is reduced, which decreases the transcriptional regulation of *Bcl-2*, *c-Myc*, and *MMP9*. While *Bcl-2* and *c-Myc* act as cancer genes by regulating cell growth, differentiation and malignant transformation, *MMP-9* is associated with tumor invasion and metastasis (46). The inhibition of the STAT3-FAK-SRC axis is implicated in lowering the cancer stem cell load, tumorigenic potential and metastasis (47). Moreover, this reduction also inhibits MMP-9 and thus tumor invasion (48) Therefore, we assessed the mRNA and protein levels in this pathway and found that the FAK-SRC-STAT axis was inhibited by cyclin G2 overexpression.

Finally, in nude mice, cyclin G2 overexpression inhibited the growth and metastasis of OSCC tumors. Because the survival of these mice is threatened by the gradual expansion of oral cancers, we used a special homemade semi-flow feeding method to ensure their survival during the experimental period. The results from the nude mice showed that cyclin G2 overexpression inhibited the growth and metastasis of OSCC. Interestingly, even though the overexpression of cyclin G2 inhibited the growth and metastasis of OSCC, it was associated with a higher grade compared to the wild type expression. These data suggest that other factors should be considered in conjunction with OSCC overexpression in order to block tumor progression. It would be worth examining human OSCC biopsies of different grades retrospectively and examining cyclin G2 expression in these cases.

This study has several limitations that should be addressed. We showed that cyclin G2, IGFBP3 and integrin show potential protein-protein interactions. However, more details are needed to understand whether these are direct interactions or if other proteins are also involved. In the future, more specific biochemical studies are needed to have a better understanding of these interactions in order to make more solid conclusions about their role in OSCC. In addition, due to the nature of OSCC in our animal model, we had to establish a special feeding method to ensure that we could maintain the mice out to 42 days. In the course of this, 1 mouse did die, and that reduced our sample size from 14 to 12. Whether this model alters the results is unknown. However, for the control mice, we observed the development of OSCC as expected. Thus, this model allowed us to test our hypothesis regarding cyclin G2 overexpression in OSCC to the best of our ability.

In summary, cyclin G2 binds to IGFBP3 and reduces the phosphorylation level of FAK and SRC and the FAK-SRC-STAT signal transduction. Our findings suggest a new mechanism of cyclin G2 for growth and metastasis in OSCC. We have illustrated the possible mechanisms (**Fig. 6**). Therefore, further study of the biological function of cyclin G2 may provide more insight into the molecular basis of the development of OSCC treatment targets.

**Figure 6.**
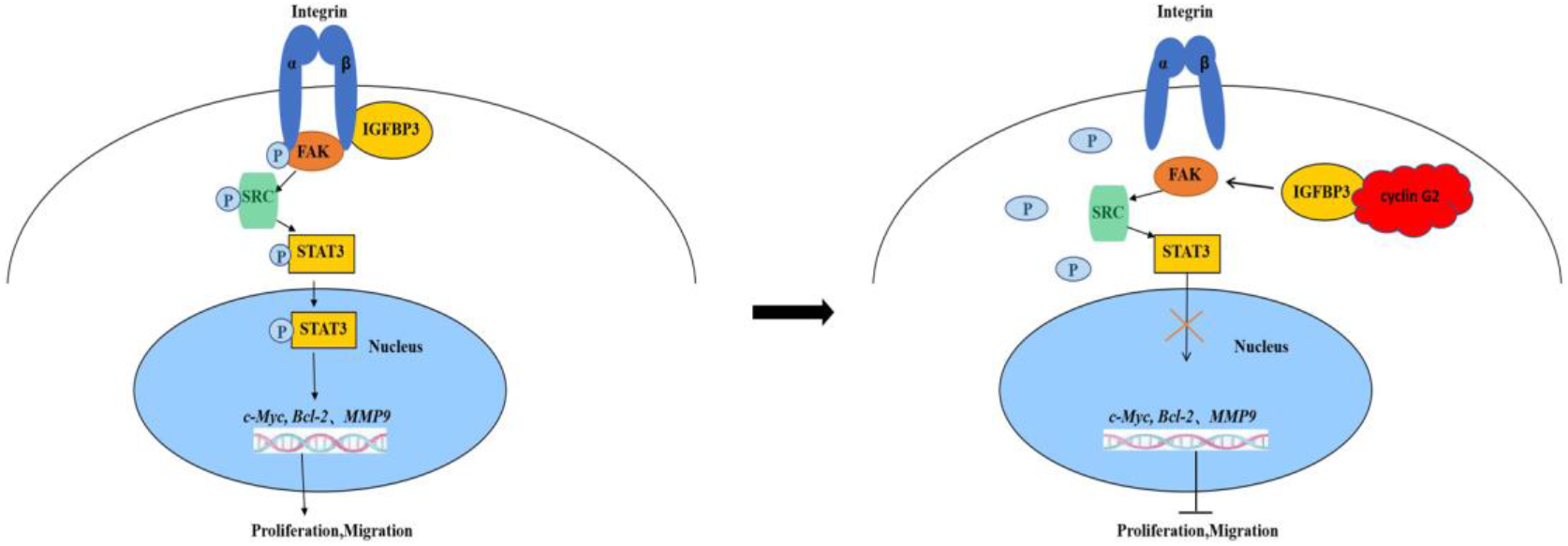
Model for cyclin G2 inhibiting the growth and metastasis of OSCC by binding to IGFBP3 and regulating the FAK-SRC-STAT signal pathway. Cyclin G2 interacts with IGFBP3 and inhibits the binding between integrin and IGFBP3 and the phosphorylation of FAK. FAK dephosphorylation inhibits the FAK-SRC-STAT signal pathway, thereby inhibiting tumorigenesis.

## Conclusions

Cyclin G2 binds to IGFBP3 and inhibits the FAK-SRC-STAT signal transduction and diminishes the growth and metastasis of OSCC. Our findings suggest a new mechanism for the role of cyclin G2 in the development of OSCC. Therefore, cyclin G2 may represent a target for OSCC treatment.

## Acknowledgments

The authors would like to thank the department of oral and maxillofacial surgery and the department of oral pathology of the oral hospital, China Medical University for providing the OSCC samples.

## Funding

This work was supported by the Foundation of the Education Department of Liaoning Province (LZDK201703, JC2019031) and the Development Program of Innovative Research Team, Ministry of Education of China (No. IRT13101).

## Availability of data and material

The datasets used and/or analyzed during the current study are available from the corresponding author upon reasonable request.

## Ethics approval and consent to participate

All the animal experiments were conducted in accordance with the guidelines of the Animal Ethics Committee of China Medical University and were approved by the Animal Ethics Committee of China Medical University.

## Patient consent for publication

This study was approved by the Research Ethics Committee of China Medical University. Informed consent was obtained from all the patients.

## Competing interests

The authors have declared that no competing interests exist.

